# Sulfatide deficiency-induced astrogliosis and myelin lipid dyshomeostasis are independent of Trem2-mediated microglial activation

**DOI:** 10.1101/2024.11.14.623651

**Authors:** Namrata Mittra, Sijia He, Hanmei Bao, Anindita Bhattacharjee, Sherry G Dodds, Jeffrey L Dupree, Xianlin Han

## Abstract

Disrupted lipid homeostasis and neuroinflammation often co-exist in neurodegenerative disorders including Alzheimer’s disease (AD). However, the intrinsic connection and causal relationship between these deficits remain elusive. Our previous studies show that the loss of sulfatide (ST), a class of myelin-enriched lipids, causes AD-like neuroinflammatory responses, cognitive impairment, bladder enlargement, as well as lipid dyshomeostasis. To better understand the relationship between neuroinflammation and lipid disruption induced by ST deficiency, we established a ST-deficient mouse model with constitutive *Trem2* knockout and studied the impact of Trem2 in regulating ST deficiency-induced microglia-mediated neuroinflammation, astrocyte activation and lipid disruption. Our study demonstrates that Trem2 regulates ST deficiency-induced microglia-mediated neuroinflammatory pathways and astrogliosis at the transcriptomic level, but not astrocyte activation at the protein level, suggesting that Trem2 is indispensable for ST deficiency-induced microglia-mediated neuroinflammation but not astrogliosis. Meanwhile, ST loss-induced lipidome disruption and free water retention were consistently observed in the absence of *Trem2*. Collectively, these results emphasize the essential role of Trem2 in mediating lipid loss-associated microglia-mediated neuroinflammation, but not both astrogliosis and myelin lipid disruption. Moreover, we demonstrated that attenuating neuroinflammation has a limited impact on brain ST loss-induced lipidome alteration or AD-like peripheral disorders. Our findings suggest that preserving lipidome and astrocyte balance may be crucial in decelerating the progression of AD.

## Introduction

Alzheimer’s disease (AD) is the main cause of dementia in the elderly population and is characterized by impairment of cognition and loss of memory (1, 2). The disease is affected by genetic, environmental, and aging factors (3, 4) such as age-related myelin disruption and loss of myelin lipids (5–7), which are important contributors to AD progression(8–10). Moreover, the myelin sheath is the foremost lesion site for the loss of various lipids in AD-like mouse models and patients (5, 9, 10). Our previous studies report that sulfatide (ST) levels were markedly lost in the white matter and the grey matter at the earliest stages of clinical AD (11). Sulfatides, also known as sulfated galactocerebrosides, are glycosphingolipids highly enriched in the brain and constitute about 10% of myelin lipids (10–13). ST regulates neuroinflammation by acting as an endogenous stimulator of the brain’s resident immune cells (9–11, 13–15). ST participates in various biological processes such as protein trafficking, cell growth, axon-myelin interactions, neuronal plasticity and cell adhesion (15). In addition, ST is an important regulator of brain lipid homeostasis (15). Studies have reported that apolipoprotein E (ApoE) functions as a modulator for ST transport (16), while the absence of *ApoE* leads to ST accumulation in mouse brain (10, 16, 17). Meanwhile, deletion of ST in mice results in major changes in brain lipid homeostasis, including severe disruption of phospholipids and sphingolipids (9–11, 13).

In addition to disrupted lipid homeostasis, enhanced neuroinflammation is another feature of AD. To understand the underlying molecular mechanism of neuroinflammation and lipid dyshomeostasis manifested in AD, our previous studies investigated the impact of ST loss using both conditional and constitutive *CST* (Cerebroside Sulfotransferase, the enzyme that catalyzes the last step of ST synthesis, encoded by *Gal3st1*) knockout mouse models. Our studies showed diverse myelin lipid disruption and neuroinflammation after ST depletion (9–11, 13, 14, 18). For example, ST deficiency induced an increase in a variety of microglial (e.g., *Trem2, Cd68, Iba-1*) and astrocytic (e.g., *Vimentin, Serpina3N, Gfap)* markers (10). Microglia, the resident macrophages in the brain and spinal cord, play a key role in AD pathogenesis by regulating neuroinflammation (19, 20). Neuroinflammatory response is one of the major defense mechanisms that promotes tissue repair by removing cellular debris and neurotoxins from the brain (19–21).These processes involve the regulation of neuronal activity which impacts disease pathogenesis (22, 23). Consequently, prolonged neuroinflammation results in neuronal cell impairment that leads to synaptic dysfunction and memory deficits in the AD brain (24).

Trem2 (triggering receptor expressed on myeloid cells 2) is one of the major receptors expressed by microglia in the brain and activates various microglial pathway genes (25). It is also highly expressed in the spinal cord (26). Evidence indicates that Trem2 plays a key role in regulating microglial responses in AD brains (10, 27). Various experimental and clinical studies have reported the activation of microglia during AD pathogenesis in aged mice as well as elderly humans (28). During AD progression, Trem2 is highly upregulated which subsequently induces the expression of various microglial genes (10, 29, 30). A disease-associated microglial (DAM) phenotype has been described for the microglial status in various neurodegenerative disease conditions (31). The transition of microglia from homeostatic to DAM is a two-stage process, where the transition from DAM stage 1 to stage 2 is Trem2 dependent. In addition to neuroinflammation, Trem2 also mediates the induction of phagocytic pathways and regulates lipid metabolism (*10, 27, 30, 32*).

Activation of astrocytes in neurodegeneration is often seen concomitantly with activation of microglia (33, 34). However, the relationship between astrogliosis and microglial activation, and the role of astrogliosis in disease progression are not very well described. Previous human and mouse transcriptomics studies showed that Trem2 regulates microglia- and astrocyte-mediated neuroinflammation at the mRNA level in AD (34). Another study reported that activated microglia can induce the activation of astrocytes at both the transcriptome and protein levels (35). In the ST-deficient mouse model, we observed an upregulated neuroinflammatory response characterized by a strong progressive activation of microglial (*Trem2*) and astrocytic (*Serpina3N*) pathway genes accompanied by lipidome disruption in the central nervous system (CNS) (10). To delineate the relationship between microglial activation and astrocytic activation, as well as lipidome disruption, we crossed constitutive *Trem2* knockout (KO) mice with conditional ST-deficient mice along with their respective controls. Interestingly, our study revealed that the presence of Trem2 is necessary to regulate the ST deficiency-induced microglial, but not astrocytic activation. Furthermore, *Trem2* does not mediate ST deficiency-induced myelin lipid imbalance and peripheral deficits (e.g., free water retention).

## Results

### *Trem2-*dependent and -independent CNS transcriptomic modulation by ST loss

In order to understand *Trem2* dependent and independent transcriptomic alterations in the CNS due to ST loss, we used NanoString gene expression analysis to assess spinal cord (SC) and cerebrum (CRM) mRNA levels in the following four groups of female and male mice (17-18 mo): *Trem2* WT/*CST* Cre- (Cre-); *Trem2* WT/*CST* Cre+ (Cre+); *Trem2* KO/*CST* Cre- (KO/Cre-) and *Trem2* KO/*CST* Cre+ (KO/Cre+) mice.

Analysis of Differentially Expressed Genes (DEGs) identified genes involved in pathways that regulate microglia, astrocytes, and immune function as shown in volcano plots in the SC of the female mice (Fig. 1A and 1B). DEGs such as *Serpina3N, Mmp12, Trem2, Cd68, Cd84, Mpeg1, Fcrls, C1qc, and ApoE* were upregulated in the *Cre*+ group compared to Cre- group, while the *Gal3st1* gene was downregulated (Fig. 1A). This indicates that ST deficiency induces microglial, astrocytic, and immune pathways at the transcriptomic level. While in the *Trem2 KO* background, only a few DEGs *(Mmp12, Clec7a, Fcrls, Serpina3N, and Ccl3*) were upregulated in *Cre+* as compared to *Cre-* (Fig. 1B). In the CRM, we detected the elevation of a few DEGs (*Serpina3N, Lilrb4a, Lcn2, Vimentin (Vim), C3, Ly9, Spp1, Ccl3, Bcl2a1*) in *Cre+* compared to *Cre-* in the *Trem2* wildtype background (Fig. S1A), while only one gene (*C4a*) was upregulated in KO/Cre*+* mice compared to *KO*/*Cre-* (Fig. S1B), indicating the changes were more robust in the SC region compared to CRM. We performed similar analysis in the male CRM, from which we only detected upregulation of one gene (Cre+ vs Cre-) (Fig. S1C) and no DEG in the *Trem2 KO* background (Cre+ vs Cre-) (Fig. S1D), suggesting discrepancies exist between sexes regarding the impact or extent of ST loss-related gene alterations. Additionally, principal component analysis (PCA) obtained from females revealed a marked separation of the Cre+ group compared to the other three groups in both SC (Fig. 1C) and CRM tissues (Fig. S1E). PCA analysis in male CRM tissue did not show much separation as compared to the females, suggesting that females are more prone to changes induced by ST loss (Fig. S1F). This is consistent with the differences between male and females during AD pathogenesis. Overall, these results indicate that *Trem2* is necessary for the induction of neuroinflammation, and immune responses triggered by ST deficiency. And the effect of ST loss is more obvious in the SC compared to CRM, as well as females compared to males.

**Fig. 1:**
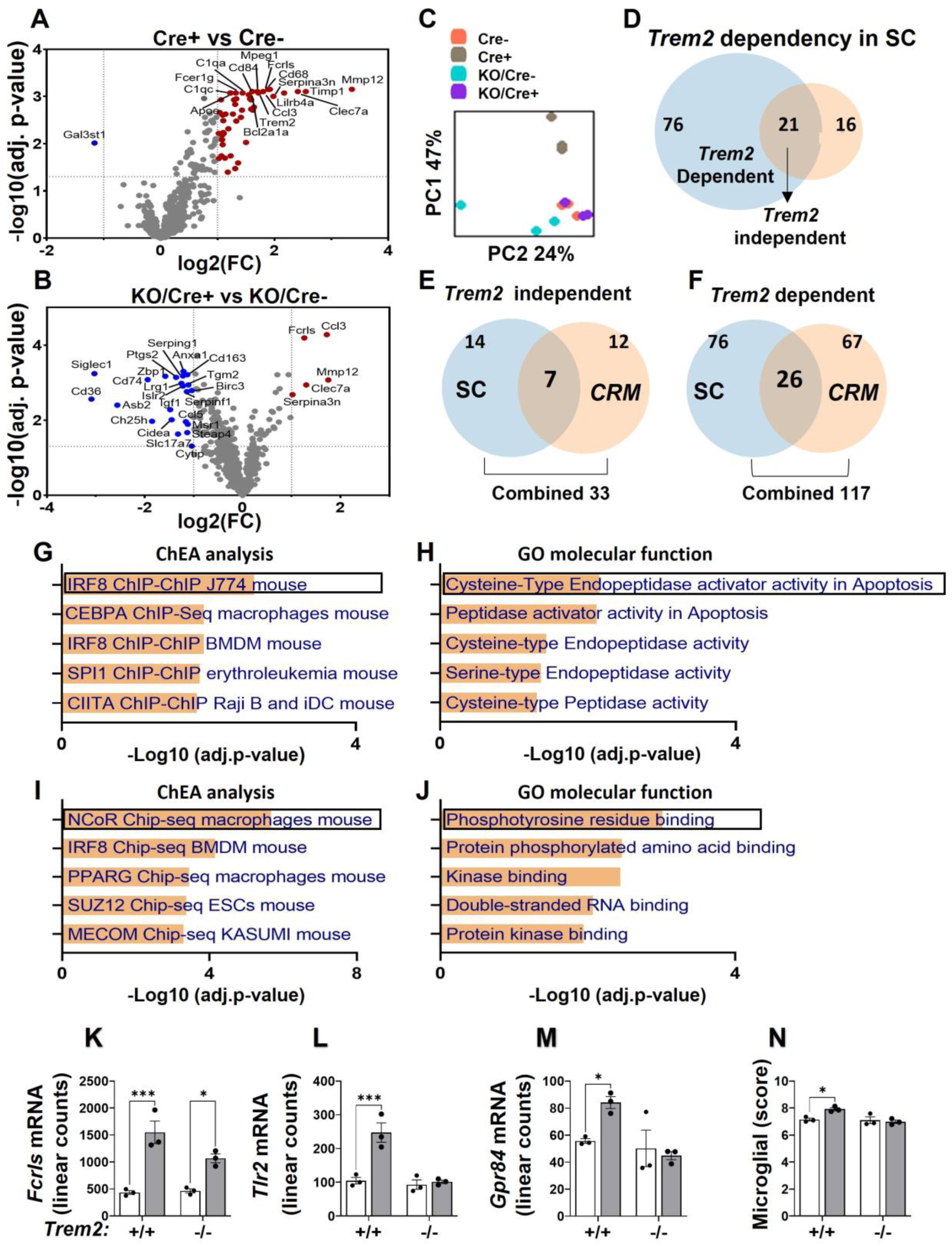
*Trem2-*dependent and -independent CNS transcriptomic modulation by ST loss. A-B. Volcano plot showing DEGs from SC (A. Female Cre+ and -, B. Female KO/Cre+ and -). C. Principal component analysis of mRNA levels obtained from the NanoString Neuroinflammatory Panel of female SC. D-F. Venn diagram showing shared DEGs that are upregulated among female genotypes or tissues. (D shared *Trem2* in SC (blue: Cre+ vs -) and (orange: KO/Cre+ vs -), E-F shared in SC and CRM, *Trem2* E. independent or F. dependent manner). G-J. Enricher analysis of shared DEGs representing TF score and ontologies based on the ChEA analysis and GO molecular function (G, H. *Trem2* independent J. *Trem2* dependent). K. mRNA linear counts of the *Trem2* independent *Fcrls* gene in female SC (white bar: Cre-, grey bar: Cre+). L-M. mRNA linear counts of *Trem2* dependent genes in female SC (L. *Tlr2*, M. *Gpr84).* N. Microglial cell scores of mRNA levels in female SC. Two-way ANOVA with multiple comparisons and a Bonferroni post-hoc test. n = 3. \**p* < 0.05, \*\**p* < 0.01, and \*\*\**p* < 0.001. Data represents the mean ± S.E.M.

To identify the *Trem2* independent/dependent transcriptomic modulations in the CNS, a Venn diagram was plotted using the upregulated DEGs from Fig. 1A and 1B (Fig. 1D). Twenty-one overlapping genes were found to be elevated by ST loss regardless of the presence of *Trem2*, suggesting the induction of these genes is *Trem2* independent (Fig. 1D). Meanwhile, the induction of 76 genes was abolished in the absence of *Trem2*; these were designated as *Trem2* dependent DEGs (Fig. 1D). Using similar criteria, we identified 19 *Trem2*-independent and 67 *Trem2* dependent DEGs from female CRM (Fig. S1G). By further referencing the gene lists between SC and CRM tissues, we found that a combined 33 DEGs were up regulated by ST loss, independent of *Trem2* across tissues, among which 7 DEGs were shared between female SC and CRM (Fig. 1E). We found a combined 117 DEGs from both SC and CRM that were *Trem2* dependent, among which 26 *Trem2-*dependent DEGs were shared by both tissues (Fig. 1F). A detailed list of 7 and 26 genes was included in the supplementary figures (Fig. S1H and S1I).

To understand the regulation and function of these genes, we analyzed the shared 7 *Trem2*-independent genes using the Enrichr database (36) and identified interferon regulatory factor 8 (*IRF8)* as the most possible up-stream transcription factor (TF) (Fig. 1G). Previous studies have reported that *IRF8* gene is involved in innate immune response (37). GO molecular functional analysis suggested that these genes may be involved in Cysteine-Type Endopeptidase activator activity (Fig. 1H), a function important for Apoptotic process (38). Similarly, we also analyzed the 26 *Trem2* dependent DEGs using Enrichr. We identified nuclear receptor corepressor (NCoR) TF as a major TF regulating these genes (Fig. 1I), and GO molecular analysis suggested these genes are mainly involved in phosphotyrosine residue and amino acid binding (Fig. 1J). NCoR is a major TF and critical regulator of macrophage inflammation and proliferation (39). *Trem2* drives the microglial response through the SYK dependent pathway, a protein tyrosine kinase which is in support of the role of *Trem2* in mediating protein kinase dependent microglia-mediated neuroinflammation (40). To confirm these findings, we investigated further details of the DEGs that were dependent/independent of *Trem2* in our datasets. Fc receptor-like molecule (Fcrls), a *Trem2* independent gene, was upregulated by ST loss in both wildtype and *Trem2* KO backgrounds in the SC and CRM from females (Fig. 1K and S1J). On the contrary, the toll like receptor 2 (*Tlr2*) and G-protein coupled receptor 84 (*Gpr84*) were only upregulated in the Cre+, but not in the KO/Cre+ compared to their respective controls in the SC (Fig. 1L and 1M) and CRM (Fig. S1K and S1L) from females. As *Trem2* is a microglia-specific gene, we evaluated the impact of ST loss and *Trem2* deletion on microglia in the CNS. The microglial cell score from females was elevated in the Cre+ compared to the Cre-, while deletion of *Trem2* abolished this induction in both SC (Fig. 1N) and CRM (Fig. S1M), suggesting deletion of *Trem2,* indeed, has greatly impaired microglial responses after ST loss.

### *Trem2* regulates ST deficiency-induced microglia-mediated neuroinflammation pathways in the CNS

Based on the above observed involvement of *Trem2* in mediating ST deficiency-induced gene transcripts, we further analyzed the impact of *Trem2* deletion in the CNS based on the mRNA levels obtained from the NanoString data. In female SC, microglial pathway genes were markedly upregulated in the *Cre*+ groups compared to those of *Cre-*, while no induction was detected in KO/Cre+ vs KO/Cre- (Fig. 2A). This indicates that the *Trem2* is necessary for activation of microglia in ST-deficiency animals. To validate this finding and provide further insight on activation status, we evaluated protein levels of microglial markers in female SC tissue by Western blot (Fig. 2B). Interestingly, we found that the absence of Trem2 ameliorated ST deficiency-induced Axl and Lilrb4a (stage 2 DAM markers) (Fig. 2C, 2D) elevation, while it had no impact on the induction of CD68 or Iba-1 (stage 1 DAM markers) (Fig. 2E, 2F). These observations were consistent with the fact that Trem2 is essential for the generation of stage 2 DAM (34).

**Fig. 2:**
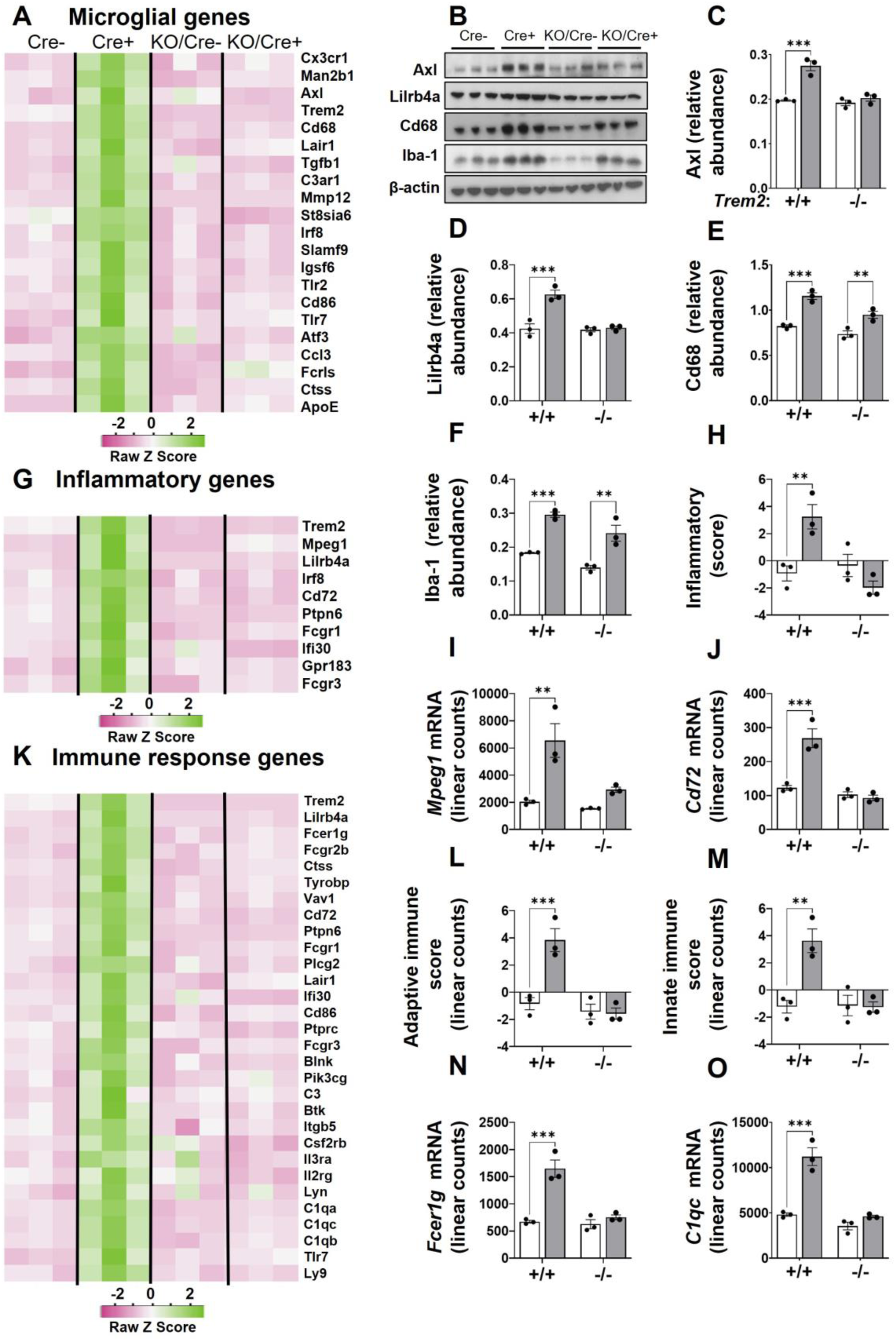
Trem2 regulates ST deficiency-induced microglia-mediated neuroinflammation and immune response pathways. A. Heatmap representing DEGs regulating the microglial genes in female SC. B. Western blots depicting protein bands of Axl, Lilrb4a, Cd68, Iba1 and β-actin in female SC. C-F Graph (white bar: Cre-, grey bar: Cre+) of each respective protein relative to β-actin in female SC (C Axl, D Lilrb4a, E Cd68, F Iba1) G. Heatmap representing DEGs regulating the inflammatory genes in female SC. H. Inflammatory pathway score in female SC. I-J. mRNA linear counts of genes in female SC (I. *Mpeg1,* J. *Cd72).* K. Heatmap representing DEGs regulating the immune response genes in female SC. L-M. Immune response pathway score in female SC (L. Adaptive, M. Innate). N-O. mRNA linear counts of genes in female SC (N. *Fcer1g,* O. *C1qc).* Two-way ANOVA with multiple comparisons and a Bonferroni post-hoc test n = 3. *p < 0.05, **p < 0.01, and ***p < 0.001. Data represents the mean ± S.E.M.

In line with the altered microglial genes, the microglial pathway score from female CRM showed an upregulation in the Cre+ group compared to Cre-, and no activation was observed after *Trem2* deletion (Fig. S2A). Similarly, the microglial genes *Cd68 and Mmp12* were elevated in the Cre+ group, not in the KO/Cre+ compared to respective controls (Fig. S2B and S2C). These results indicate that *Trem2* mediates ST deficiency-induced microglial activation at the transcriptome level specifically via promoting the formation of stage 2 DAM.

Enhanced inflammatory and immune pathways were detected in ST deficiency based on our previous study (10), yet it is unknown whether these effects are mediated by *Trem2*. By analyzing the levels of a group of inflammatory pathway genes, including *Fcgr1, Ptpn6, Ifi30, Fcgr3, Lilrb4a, Gpr183, Cd72, Irf8, Mpeg1,* and *Trem2*, we found that the induction of inflammation by ST loss was no longer observed in a *Trem2* KO background (Fig. 2G). Accordingly, deletion of *Trem2* abolished the upregulation of the inflammatory pathway score (Fig. 2H), as also reflected by the gene levels of *Mpeg, Cd72*, (Fig. 2I-J) and *Trem2* (Fig. S2D). Similar results were found in the female CRM, where the induction of pathway score (Fig. S2E) and inflammatory genes (*Mpeg, Lilrb4a*) (Fig. S2F and S2G) were upregulated in the Cre+ mice only in the presence of *Trem2*.

### *Trem2* regulates ST deficiency-induced adaptive and innate immune response pathways

To understand the role of *Trem2* in the ST deficiency-induced adaptive and innate immune response pathways, we assessed immune-related genes in female SC and observed a clear upregulation in the Cre+ group compared to that of Cre-mice, but no induction was detected in the *Trem2* KO background (Cre+ vs Cre-) (Fig. 2K). Consistently, both adaptive and innate immune response pathway scores were upregulated in the Cre+, while the induction was diminished in KO/Cre+ compared to KO/Cre- (Fig. 2L and 2M). The upregulation of immune response genes such as *Fcerg1* and *C1qc* by ST loss was no longer detected in the Cre+ vs Cre- in the *Trem2* KO background (Fig. 2N and 2O). Similar trends of adaptive, innate immune response pathway scores (Fig. S2H and S2I) and mRNA linear counts of the *C1qa* and *C1qc* genes (Fig. S2J and S2K) were observed in the female CRM. Notably, the inflammatory, immune and microglial pathways in male CRM also showed a weakened induction by ST loss in the absence of *Trem2* (Fig. S2L). Conclusively, *Trem2* is necessary for activating the ST deficiency-induced neuroinflammation pathway and immune response at the transcriptomic level in both female SC and CRM, while male CRM showed minimal effects, suggesting the female effect to be more robust.

### *Trem2* regulates ST deficiency-induced astrocyte activation at the transcriptomic level but not at the protein level in the CNS

We next assessed the impact of *Trem2* deletion on astrocyte activation induced by ST deficiency. A group of genes that are associated with astrocyte function was evaluated based on results from our NanoString measurements. We found that in female SC, ST deficiency-induced astrocyte activation-related genes such as *Cd109, Vimentin (Vim), Tm4sf1, Serpina3N, Ptx3, S1pr3, Cd14, Timp1, ApoE,* and *Osmr* in Cre+ compared to Cre-, while knockout of *Trem2* greatly limited these inductions, as shown in Fig. 3A. In line with this, the astrocyte pathway scores and mRNA levels of *ApoE, Serpina3N*, and *C4a* were found upregulated in Cre+ vs Cre- only in the presence of *Trem2* (Fig. 3B-E). A similar trend was also observed in the female CRM, as shown in Fig. S3A-E. In male CRM, we detected a pattern resembling that of females, but the regulation was less robust when compared to females (Fig. S3F and S3G).

**Fig. 3:**
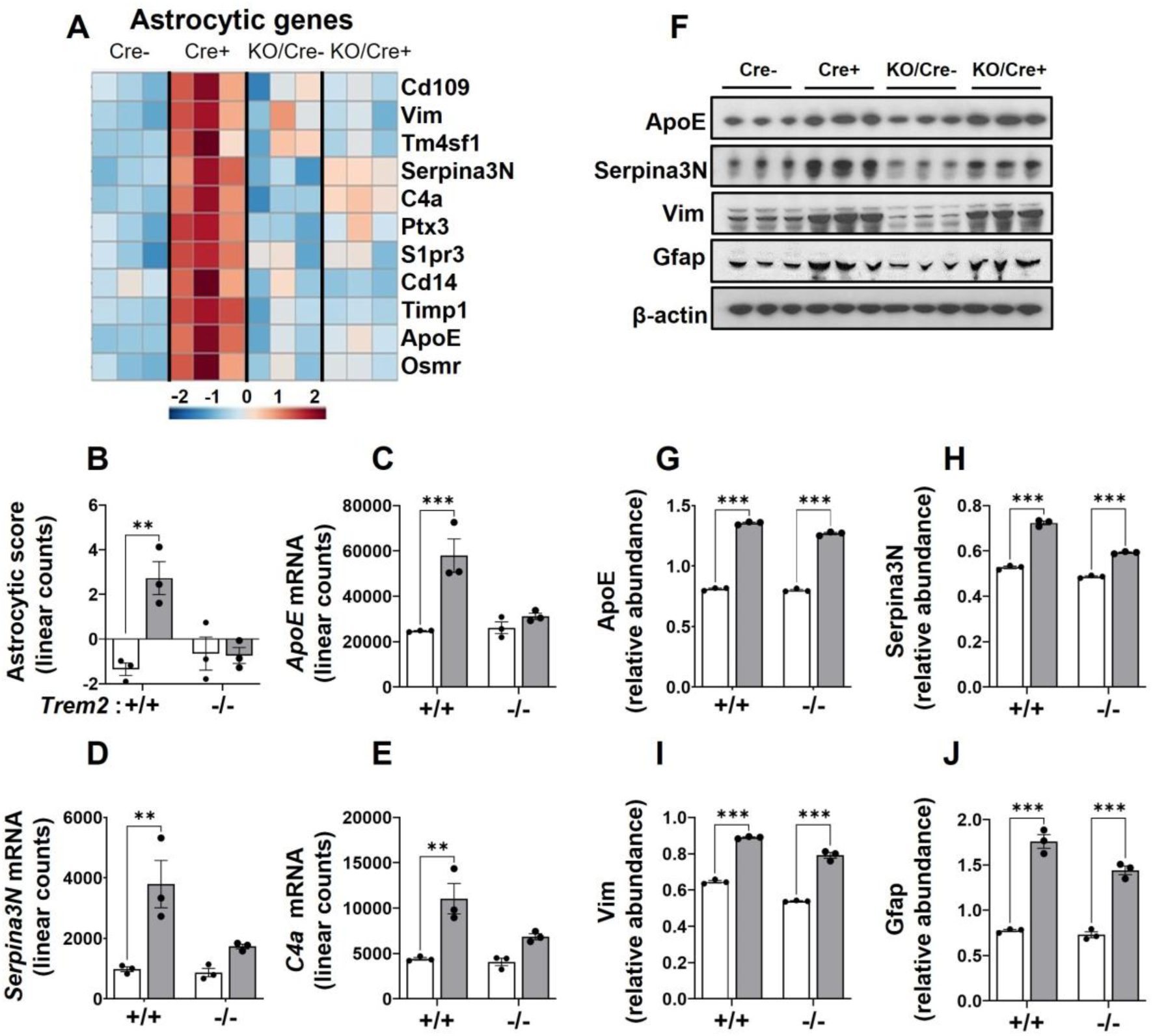
*Trem2* regulates ST deficiency-induced astrocyte activation at the transcriptomics level, but not at the protein level. A. Heatmap from DEGs regulating the astrocytic genes in female SC. B. Astrocytic score in female SC (white bar: Cre-, grey bar: Cre+). C-E. mRNA linear counts of genes in female SC (C. *ApoE,* D. *Serpina3N,* E. *C4a*). F. Western blots depicting protein bands of ApoE, Serpina3N, Vim, Gfap and β- actin in female SC. G-J. Graph (white bar: Cre-, grey bar: Cre+) of each respective protein relative to β-actin in female SC (G ApoE, H Serpina3N, I Vimentin, J Gfap). Two-way ANOVA with multiple comparisons and a Bonferroni post-hoc test n = 3. *p < 0.05, **p < 0.01, and ***p < 0.001. Data represents the mean ± S.E.M.

To further validate our findings from the transcription level, we assessed protein levels of astrocytic markers in the female SC tissues using Western blots. Surprisingly, in contrary to mRNA levels, the protein levels of ApoE, serpina3N, Vim, and GFAP were markedly upregulated in the Cre+ group compared to the Cre-group regardless of the presence or absence of Trem2 (Fig. 3F-J). Collectively, these results suggest that Trem2 regulates the ST deficiency-induced astrocyte activation at the mRNA level, but not at the protein level in SC.

### Sulfatide deficiency-induced alterations in myelin lipids are independent of *Trem2*

Along with microglia and astrocyte activation, significant disruption of the lipidome was observed in both CRM and SC of ST-deficient mice (10, 14). To determine if *Trem2* plays a role in ST deficiency-induced lipid disruption, we performed multi-dimensional mass spectrometry-based shotgun lipidomic analysis on both female and male CNS.

In female CRM, lipidomic data showed that ST deficiency-induced losses of myelin lipids, e.g., phosphatidic acid (PA), cerebroside (CBS) and acyl carnitine (AC), were significantly reduced in both Cre+ vs Cre- (indicated by *) and KO/Cre+ vs KO/Cre- (indicated by $) (Fig. 4A). Detailed group-wise comparisons of ST, PA, CBS, and plasmalogen attenuation in both *Cre+* and KO/Cre+ groups compared to their respective controls were shown in Fig. 4B-E. The results indicate that ST deficiency-induced myelin lipid disruption is independent of *Trem2*. Meanwhile, the ST loss-driven alterations in the cardiolipin (CL) and phosphatidylinositol (PI) were only present in the Cre+ vs Cre-, not in the (KO/Cre+ vs KO/Cre-), indicating these changes to be *Trem2* dependent (Fig. 4F and 4G).

**Fig. 4:**
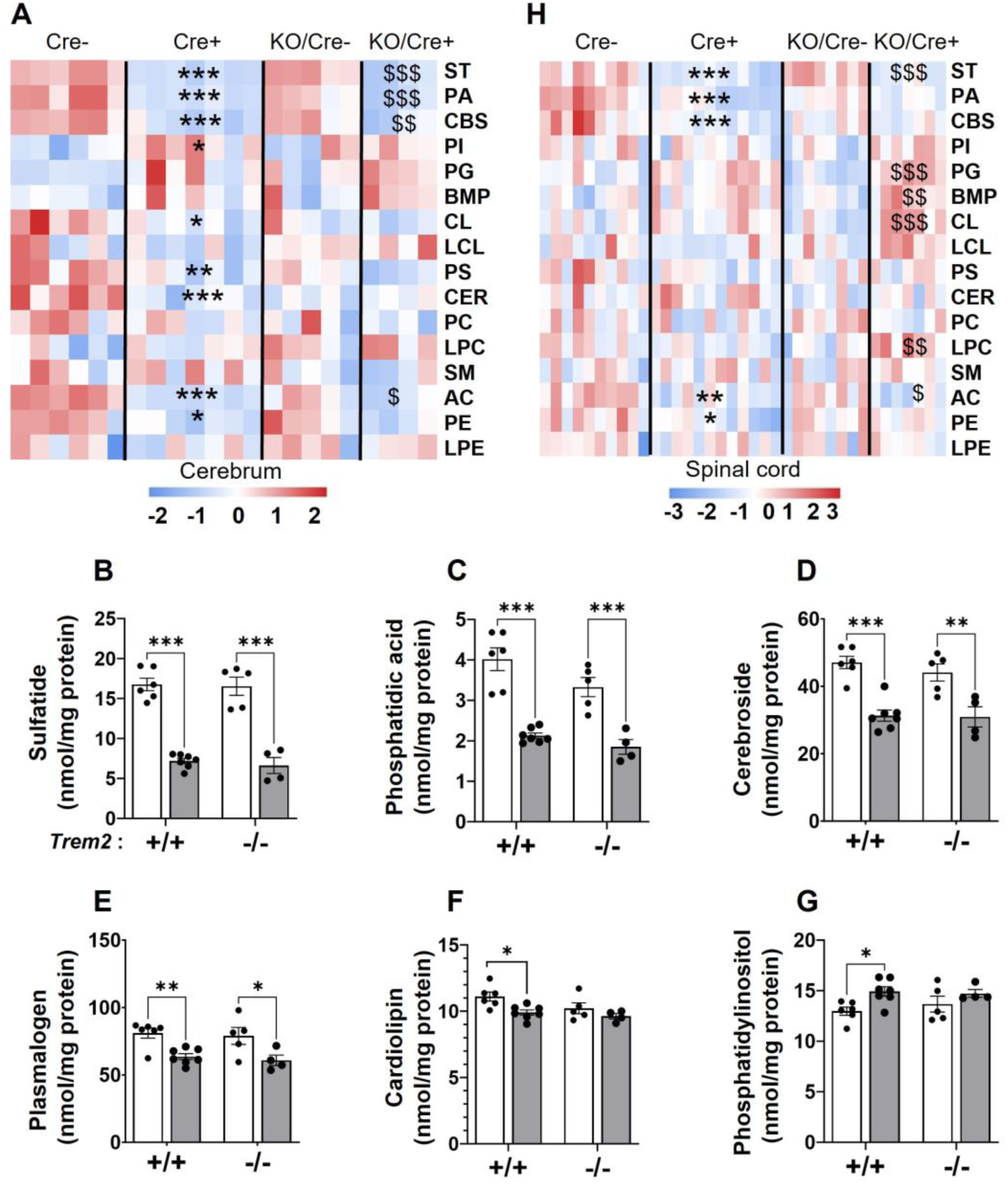
Sulfatide deficiency-induced alterations in myelin lipids are independent of *Trem2.* A. Heatmap of total lipid classes from lipidomics data in female CRM (* indicates Cre*+* vs Cre- and $ indicates KO/Cre+ vs KO/Cre-). B-G. Graph of each respective lipid (white bar: Cre-, grey bar: Cre+) in female CRM (B. ST, C. PA, D. CBS, E. Plasmalogen, F. CL, G. PI). H. Heatmap of total lipid classes from lipidomics data in female SC (* indicates *Cre*+ vs *Cre*- and $ indicates KO/Cre+ vs KO/Cre-). Two-way ANOVA with multiple comparisons and a Bonferroni post-hoc test n = 5-14. *p < 0.05, **p < 0.01, and ***p < 0.001. Data represents the mean ± S.E.M.

In female SC, ST deficiency induced the reduction of mitochondria-specific AC species in both Cre+ (indicated by *) and KO/Cre+ (indicated by $) groups compared to their respective controls (Fig. 4H). Interestingly, other classes of lipids such as phosphatidylglycerol (PG), Bis(monoacylglycero)phosphate (BMP), cardiolipin (CL), and lysophosphatidylcholine (LPC) were elevated only in the KO/Cre+ group, indicating the above alterations were specifically responsive to ST loss only when *Trem2* is absent. Comparisons of ST, PA, CBS, and PG molecular species between groups are summarized in supplementary materials (Fig. S4A-D). In male SC, ST deficiency led to a reduction of myelin lipids (e.g., PA, PS, and PE), and these alterations were independent of *Trem2* (Fig. S4E). Group-wise comparisons are shown in Fig. S4F and S4G.

Overall, our lipidomic measurements suggested that ST deficiency-induced alterations in myelin lipids are mostly independent of *Trem2* in both sexes.

### Sulfatide deficiency-induced free water retention in females is independent of *Trem2*

We recently reported that prolonged ST loss leads to bladder enlargement due to elevated urine retention, reflecting impaired CNS control of peripheral organ function in AD (14). The elevated urine phenotype was measured using qMRI and displayed as free water retention. To access if *Trem2* is involved in the regulation of ST deficiency-induced free water retention, we measured the free water weight in 12- and 17-mo female and male mice from all groups using quantitative MRI (qMRI). Significant elevation in free water retention was observed in 17-mo-female mice but not at 12 mo in the Cre+ and KO/Cre+ mice compared to their respective controls (Fig. 5A-B). No significant difference in free water was detected in males at 17 or 12 mo (Fig. 5C-D). Note that in addition to free water, we also evaluated other parameters, including total water weight (Fig. 5E-H), lean weight (Fig. S5A-D), fat weight (Fig. S5E-H), and body (Fig. S5I-L) weight. No change was observed among all groups at the age of 12 and 17 mo. These results suggest that ST deficiency-induced urine retention is independent of *Trem2*.

**Fig. 5:**
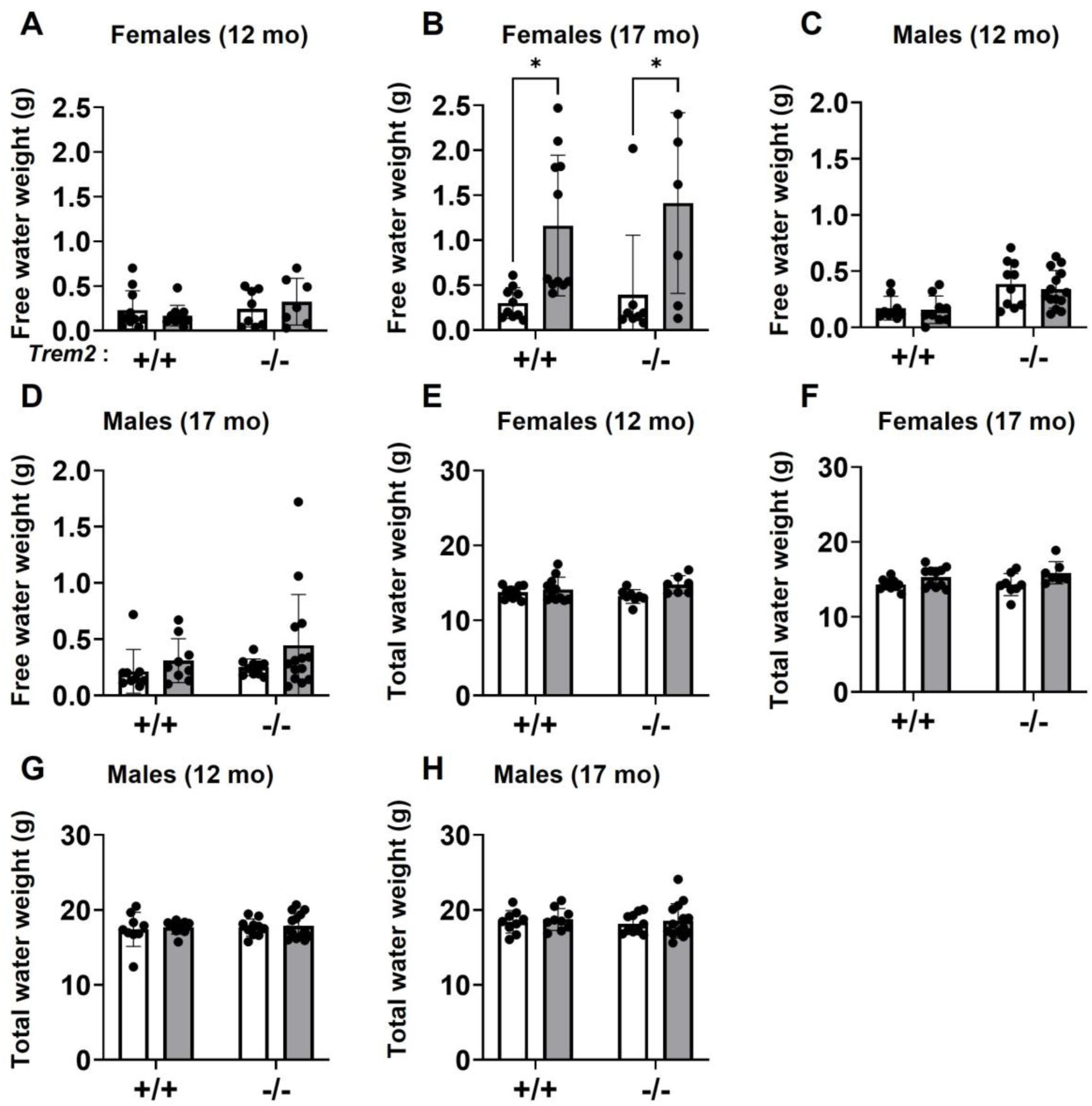
Sulfatide deficiency-induced free water retention in female animals is independent of *Trem2*. A-D. Free water weight (A. 12-mo females, B. 17-mo females, C. 12-mo males, D. 17-mo males). E-H. Total water weight (E. 12-mo females, F. 17-mo females, G. 12-mo males, H. 17-mo males). All graphs (white bar: Cre-, grey bar: Cre+). Two-way ANOVA with multiple comparisons. n = 5-14. *p < 0.05, **p < 0.01, and ***p < 0.001. Data represents the mean ± S.E.M.

## Discussion

Our previous studies have shown that ST is decreased in AD pathogenesis (10). ST reduction induces neuroinflammation and disrupts the brain and spinal cord lipidomes leading to cognitive impairment, as well as bladder enlargement (9, 10, 12, 14). Trem2 has been shown to be an essential factor in mediating microglial activation (25) and is induced in CNS tissue of ST-deficient mouse (10). Interestingly, previous studies reported that *Trem2* also *a*ffects the lipid metabolism of myelin, impacting phospholipids and cholesterol, and promotes the transition of homeostatic microglia into a disease-associated microglia phenotype (41). Based on this evidence, the current study aims to investigate the role of Trem2 in ST deficiency-induced microglia-mediated neuroinflammation and astrocyte activation, as well as lipid disruption in the CNS.

To achieve this goal, we crossed our established conditional myelin sulfatide deficiency mouse model (CST KO) with a constitutive *Trem2* KO mouse line and administered tamoxifen at 3 mo of age to induce ST loss post myelin maturation. The ST-deficient mouse model on a wild type background mimics the ST loss that occurs in AD brains leading to cognitive impairment, neuroinflammation, and lipid disruption (24–26). Using NanoString nCounter gene expression analysis, we further explored the effects of ST deficiency on mRNA levels in the presence and the absence of *Trem2*. Based on previous publications, *Trem2* is essential for stage 2 activation of DAM signature (34), while *Trem2* KO downregulates the transcription of microglial markers and complement components, resulting in attenuation of microglial density and phagocytosis (42). In our study, ST loss in the wild type background led to profound upregulation of DEGs that regulates different microglia-associated biological pathways (such as *C1qa, Mpeg1, Cd84, C1qc, ApoE, Trem2, Mmp12, Fcerg1g, Fcrls, Cd68, Timp1, Lilrb4a, Ccl3, Clec7a* and *Bcl2a1a*), while the induction of these genes was almost completely abolished (except for *Ccl3, Mmp12, Clec7a, Fcrls, and Serpina3N*) in the *Trem2* KO background. This suggests that *Trem2* is critical for the activation of microglia in ST deficiency. Further, by validating protein levels of microglial markers in our mouse models, we identified that *Trem2* KO did not impact the induction of stage 1 DAM (e.g., CD68 and Iba-1), but ameliorated the induction of stage 2 DAM markers (e.g., Axl, Lilrb4a) in ST deficiency. This is in line with the fact that *Trem2* functions as a “gate keeper” of stage 1 to stage 2 DAM transition (34). Consistent with the blunted microglial activities in *Trem2* KO mice (both Cre+ and Cre-), both inflammatory and immune response pathways were silenced by *Trem2* deletion despite the loss of ST, emphasizing the fact that ST loss-induced immune response and neuroinflammation largely depend on *Trem2*.

In addition to *Trem2*-dependent genes, we also identified a group of genes that were upregulated in ST deficiency independent of *Trem2*, including *Ifnar1, Pecam1, Csf1r, Fcrls, Asb2, Itgam*, and *Csf3r*. Interestingly, most of these genes are highly enriched in microglia (except for *Ifnar1* and *Pecam1*, which are mostly expressed in endothelial cells), suggesting an additional mechanism governing microglial response in the absence of *Trem2*. In line with this, a recent study highlighted the involvement of Trem2-independent microgliosis in a neurodegeneration model characterized by tau pathology (43). Specifically, a group of microglial lysosomal genes, including *Ctsd*, *Ctsb*, and *Cd68*, were upregulated in tau pathology independent of Trem2 (43). Notably, our findings also revealed a persistent elevation of Cd68 levels in ST deficiency in the absence of Trem2, suggesting that microglial lysosomal activation may be one of the Trem2-independent mechanisms mediating microglial responses in ST deficiency

One of the key questions we ask in this study is whether microglial activation plays an essential role in ST loss-induced phenotypes (including astrocyte activation and lipidome disruption, as well as peripheral phenotype such as urine retention). Activation of both microglia and astrocytes is well-described in AD and is consistently detected in our ST-loss models. It is shown that astrocytic activation in AD promotes proteostatic, inflammatory, and metal ion homeostasis pathways, while microglial activation leads to enrichment in phagocytosis, inflammation, and proteostasis pathways (44). The existence of common and cell-specific responses between these two cell types indicates functional similarity and diversity. It has been suggested that reactive astrocytes are induced by activated microglia (35, 45). Conversely, other studies showed that astrocytes mediate microglial activation through C3-C3aR complement signaling (46). By deletion of *Trem2*, we eliminated microglial activation in our mouse models. The observation that several reactive astrocytic markers remain elevated in the *Trem2* KO background suggests that ST deficiency induces astrocyte activation independent of microglia-mediated immune and inflammatory responses. Given that astrocytes interact with both neural and non-neural cells (including neurons and their synapses, oligodendrocytes, oligodendrocyte progenitor cells, microglia, various perivascular cells, meningeal fibroblasts, and circulating immune cells) (47), it is possible that myelin or neuronal alterations triggered by ST loss may contribute to the activation of astrocytes in our mouse models. Further investigation is needed to elucidate the triggering factor for astrogliosis in ST deficiency.

In addition to the ST-induced effects from our previous studies, ST deficiency is responsible for triggering major myelin lipid disruption and bladder enlargement (9, 10, 14, 15). We and others have previously established that ST deficiency disrupts the myelin lipid network of a variety of phospholipids and leads to axonal degeneration (48). Meanwhile, *Trem2* can impact *ApoE*, which participates in lipid transport as a major lipid carrier (49, 50). During our investigation, we detected ST deficiency-induced myelin lipid disruption regardless of the presence or absence of *Trem2*, particularly in CRM. This indicates that Trem2-dependent microglia-initiated neuroinflammation does not contribute to the lipidome disruption induced by ST deficiency.

Chronic CNS ST loss leads to bladder enlargement in mice, which represents the loss of function in controlling peripheral organs at the late stages of AD. Although neuroinflammation was believed to be a mediator of such phenotype, no previous studies have demonstrated the extent of its contribution. Here we utilized our novel mouse model of ST loss with neuroinflammation elimination and demonstrated that microglia-mediated neuroinflammation is not essential for bladder enlargement in ST deficiency. This strongly suggests that astrocyte activation or disruption of the lipidome is likely the actual main mediator(s) of impaired (CNS or peripheral) control in late AD.

In line with the fact that females are at a higher risk for developing AD than males, the neuroinflammation, lipidome disruption, and bladder phenotypes were consistently more severe in females than males in our current and previous studies (9, 10, 14). Several reasons may be at play for this outcome: the difference in the extent of ST loss between sexes may lead to a different degree of response by cells in the CNS; glia or neurons in females could be more sensitive to environmental lipid changes; and genetic and hormonal discrepancies between sexes can affect the susceptibility of CNS to disease triggering factors. Although the exact causes of sex differences have not been demonstrated, we included studies for both sexes to obtain more comprehensive observations on both functional and mechanistic aspects.

Although AD is considered a brain disorder, the potential role of SC in neurodegeneration has gained increased recognition (51, 52). Through investigating both SC and CRM tissues in the current study, we demonstrated a significant overlap of transcriptomic and functional responses between these two CNS regions, suggesting the existence of common regulatory mechanisms. On the other hand, several region-specific DEG groups were identified between SC and CRM. Whether these signatures reflect region-specific function will require further investigation. An additional observation is that in evaluating the astrocyte activation status, we observed discrepancies between transcriptional and protein levels. This suggests involvement of post-translational mechanisms in regulating astrocyte activation. It also emphasizes the importance of evaluating both mRNA and protein levels in determining the contribution of a molecule of interest.

In summary, we have established the critical role of Trem2 in mediating microglia activation within a model of ST loss that closely resembles the AD phenotype. Our findings demonstrate that while Trem2 is essential for microglia-driven neuroinflammation, it has a limited impact on astrogliosis or lipidome alterations associated with myelin ST deficiency (Fig. 6). These conclusions provide new insights into glia-glia interactions and may contribute to the understanding of neuroinflammatory mechanisms in AD.

**Fig. 6:**
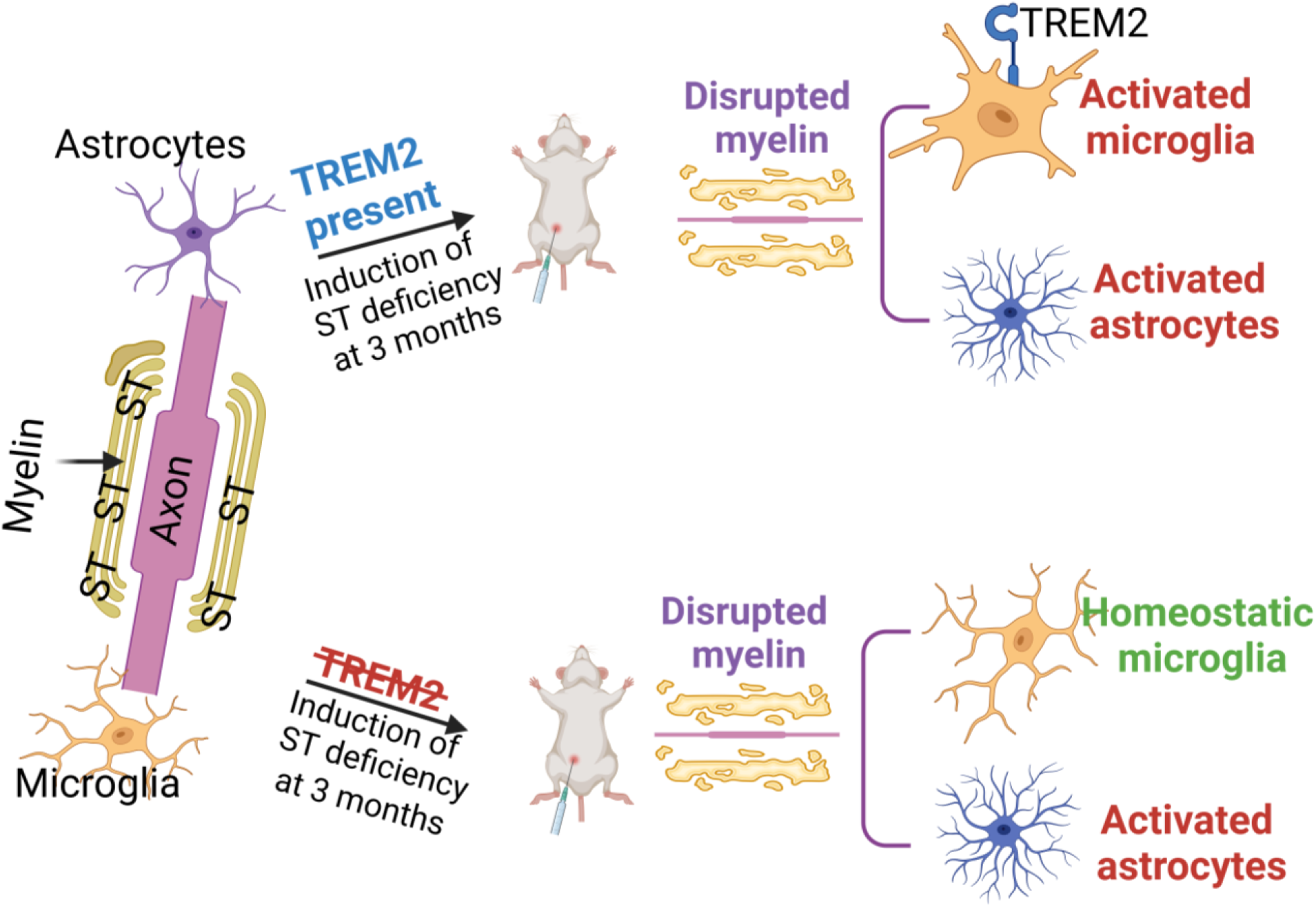
Role of *Trem2* in ST deficiency-induced neuroinflammation and lipid disruption. Schematic summary showing the necessity of *Trem2* in regulating ST deficiency-induced microglia-mediated neuroinflammation specifically, but not astrocyte activation or myelin lipid dyshomeostasis.

## Materials and Methods

### Mice

The *CST* fl/fl mice were generated by Applied Stem Cell, Inc. using CRISPR technology by injecting C57BL/6 embryos as described previously (10, 48). The Plp1-*Cre*ERT and *Trem2* KO mouse lines were both purchased from JAX (stock #005975 and #027197, respectively). The *CST* fl/fl mice were crossed with Plp1-*Cre*ERT+ mice to generate conditional ST-deficient (Cre+) mice along with their respective (Cre-) controls as previously described (10). The constitutive *Trem2* KO mice were crossed with ST-deficient Cre+ mice to generate *Trem2* KO ST-deficient mice (*Trem2, CST* fl/fl/Plp1- *Cre*ERT) denoted as (KO/Cre+) along with the control (KO/Cre-).

Tamoxifen (40 mg/kg body weight, Sigma, Cat. #: T5648-5G) was administered via intraperitoneal injections once every 24 h for 4 consecutive days to male and female control *Trem2* WT/*CST* Cre- (Cre-); *Trem2* WT/*CST* Cre+ (Cre+); *Trem2* KO/*CST* Cre- (KO/Cre-) and *Trem2* KO/*CST* Cre+ (KO/Cre+) mice at 3 mo of age. Mice were housed in groups of ≤5 mice/cage in a temperature and humidity-controlled environment under a 12-h light /12-h dark cycle. They were provided with food and water *ad libitum*. The protocols for animal experiments were conducted in accordance with the ‘Guide for the Care and Use of Laboratory Animals’ (8^th^ edition, National Research Council of the National Academies, 2011) and were approved by the Institutional Animal Care and Use Committee at University of Texas Health San Antonio.

### Body Composition

Quantitative Magnetic Resonance Imaging (qMRI) was used to assess the body composition (fat weight, lean weight, free water, and total water) of individual mice using an EchoMRI 3-in-1 Body Composition Analyzer (Echo Medical Systems, Houston, TX, USA).

### Brain preparation

For biochemical analysis, animals were anesthetized, cerebrum (from the left hemi-brain) and spinal cord tissues were collected, and flash frozen in liquid nitrogen.

### Gene expression analysis

Cerebrum and spinal cord tissues were powdered in liquid nitrogen followed by RNA extraction using the Animal Tissue RNA Purification Kit (Norgen, Canada, Cat. #: 25700) according to the manufacturer’s protocol. The concentration of RNA was determined using a DS-11 Spectrophotometer (DeNovix, USA). NanoString multiplex gene expression analysis was performed using the Mouse Neuroinflammation Panel on the nCounter® Max machine Profiler according to the manufacturer’s protocol (NanoString Technologies, USA). Data were analyzed using nSolver 4.0 software. Our basic analysis included the default Positive Control Normalization Parameters which include a geometric mean calculation and flags lanes if their normalization factor is outside of the 0.3-3.0 range. We also set the Code Set Content to standard fold change estimation for all pairwise ratios. Our advanced analysis included custom analysis with all default settings except selecting P-value adjustments (Benjamini Hochberg) and (use all probes) under the section of differential profiling and cell type profiling.

### Lipid extraction and mass spectrometric analysis of lipids

Multidimensional mass spectrometry-based shotgun lipidomics was performed as previously described (13). Briefly, pulverized frozen CRM and SC tissues were homogenized in ice-cold phosphate-buffered saline using a Precellys® Evolution Tissue Homogenizer (Bertin, France). The protein concentration of homogenates was separately determined with a Pierce BCA Protein assay (Thermo Fisher, USA, Cat# 23225) according to the manufacturer’s instructions. Lipids were extracted by the modified procedure of Bligh and Dyer extraction in the presence of internal standards that were added based on the total protein content of each sample (53). Lipids were quantified by ion peak intensity comparison to the internal standard of the class of lipids as acquired by a triple-quadrupole mass spectrometer (TSQ Altis, Thermo Fisher Scientific, Waltham, MA, USA) equipped with a TriVersa NanoMate® device (Advion Interchim Scientific, USA) on an Xcalibur operating system as previously described (54). Data processing including ion peak selection, baseline correction, data transfer, peak intensity comparison and quantitation was performed as previously described (10, 55). The results were normalized to the protein content (nmol lipid/mg protein).

### Western blotting

Frozen spinal cord tissue was powdered in liquid nitrogen, followed by homogenization in N-PER™ Neuronal Protein Extraction Reagent (Thermo Scientific, USA, Cat. #: 87792) with Halt Protease and Phosphatase Inhibitor Cocktails (Thermo Scientific, USA, Cat. #: 78430) using a Precellys® Evolution Tissue Homogenizer (Bertin, France). Homogenates were cleared by centrifugation at 12000 x g for 30 min at 4 °C. Supernatants were collected and their protein concentrations were determined using the Pierce^TM^ BCA Protein Assay Kit (Thermo Scientific, USA, Cat. #: 23225). Forty micrograms of protein per sample were separated on NuPAGE 4–12%, Bis-Tris gels (Invitrogen, USA, Cat. #: NP0323) under reducing conditions. Separated proteins were then transferred to PVDF (Fluor) membrane (Millipore, USA, Cat. #: IPVH00010) for (1 h at 100 V) using NuPAGE transfer buffer (Invitrogen, USA, Cat. #: NP00061). Membranes were hybridized overnight at 4°C using a 1:1000-1:2000 dilution of the following primary antibodies: anti-GFAP (rabbit, Thermo Fisher, USA, Cat. #: PA1-10019), anti-CD68 (rabbit, Thermo Fisher, USA, Cat. #: 97778S); anti-ApoE (rabbit, Abcam, USA, Cat. #: NBP1-49529SS), anti-Serpina3N (goat, R&D systems, USA, Cat. #: AF4709), anti-Vimentin (chicken, Abcam, USA, Cat. #: ab24525), anti-Iba1 (rabbit, Fujifilm Wako, USA, Cat. #: 016-20001), anti-Axl (goat, R&D systems, USA, Cat. #: AF854), and anti-β-actin (rabbit, Cell Signaling Technology, USA, Cat. #: 3700S). All primary antibodies were followed by hybridization for 1h at room temperature with a 1:3000 dilution of the appropriate horseradish peroxidase (HRP)-linked secondary antibody: goat anti-rabbit (Cell Signaling Technology, USA, Cat. #: 7074), donkey anti-goat (Thermo, USA, Cat. #: PA1-28664), or goat anti-chicken (Thermo Scientific, USA, Cat. #: A-16054). Membranes were washed 3 x 10 min in tris buffered saline buffer after each antibody incubation. Pierce™ SuperSignal™ West Atto Ultimate Sensitivity ECL Substrate (Thermo Scientific, USA, Cat. #: A38556) was used to detect proteins before exposure to autoradiography film (HyBlot CL, Denville Scientific). Some membranes were reblotted after being incubated in protein stripping buffer (Thermo Fisher Scientific, USA, Cat. #: 46430). The film was scanned, and protein expression was analyzed using Alpha Imager Software (Alpha Innotech). All protein expressions were normalized to β-actin.

### Statistics

Data were presented as mean ± SEM. All statistical analyses for the NanoString data were performed using NanoString nSolver 4.0 software according to the manufacturer’s recommendations. All other statistical analyses were performed using Prism 9 (GraphPad). Two-way ANOVA with Bonferroni post-hoc test was conducted for multiple comparisons. \**p* < 0.05, \*\**p* < 0.01, and \*\*\**p* < 0.001 were used to compare the mean values across multiple groups

## Abbreviations

AC: Acyl carnitine
AD: Alzheimer’s disease
Apo: Apolipoprotein
BMP: Bis(monoacylglycero)phosphate
CBS: Cerebroside
CER: Ceramide
CL: Cardiolipin
CNS: Central nervous system
CD68: Cluster of differentiation 68
CD33: Cluster of differentiation 33
DAM: Disease associated microglia
DEGs: Differentially expressed genes
GSEA: Gene set enrichment analysis
LPC: Lysophosphatidylcholine
LPE: Lysophosphatidylethanolamine
LCL: Lysocardiolipin
PA: Phosphatidic acid
PC: Phosphatidylcholine
PE: Phosphatidylethanolamine
PG: Phosphatidylglycerol
PI: Phosphatidylinositol
PS: Phosphatidylserine
GFAP: Glial fibrillary acidic protein
Iba-1: Ionized calcium binding adaptor molecule-1
IRF8: Interferon regulatory factor 8 SM Sphingomyelin
ST: Sulfatide
TNF: Tumor necrosis factor
TF: Transcription factor
MAPK: Mitogen activated protein kinase
JNK: c-Jun N-terminal Kinase
NFκB: Nuclear Factor Kappa B
Mmp12: Matrix Metallopeptidase 12
PLS-DA: Partial Least Squares-Discriminant Analysis

## Acknowledgements

We would like to thank the University of Texas at San Antonio Genomics Core for performing NanoString multiplex gene expression analysis, and the San Antonio Nathan Shock Center for performing the qMRI body composition analysis.

## Contributions

SH and XH were involved in the initial conceptualization of the project. NM and SH designed, performed experiments and analyzed results. HB, SD, and AB contributed to animal handling and data measurements. JLD contributed to the generation of mouse models and discussion of the project. SH and NM wrote the manuscript. SD and XH edited the manuscript. XH directed the project and provided laboratory resources.

## Ethics approval and consent to participate

The animal protocol was approved by the Institutional Review Board of UT Health San Antonio (protocol code 20180044AP and approved on 01-09-2019).

## Consent for publication

All authors have approved the contents of this manuscript and provided consent for publication.

