## Supplemental materials for "Sulfatide deficiency-induced astrogliosis and myelin lipid dyshomeostasis are independent of Trem2-mediated microglial activation"

Fig. S1

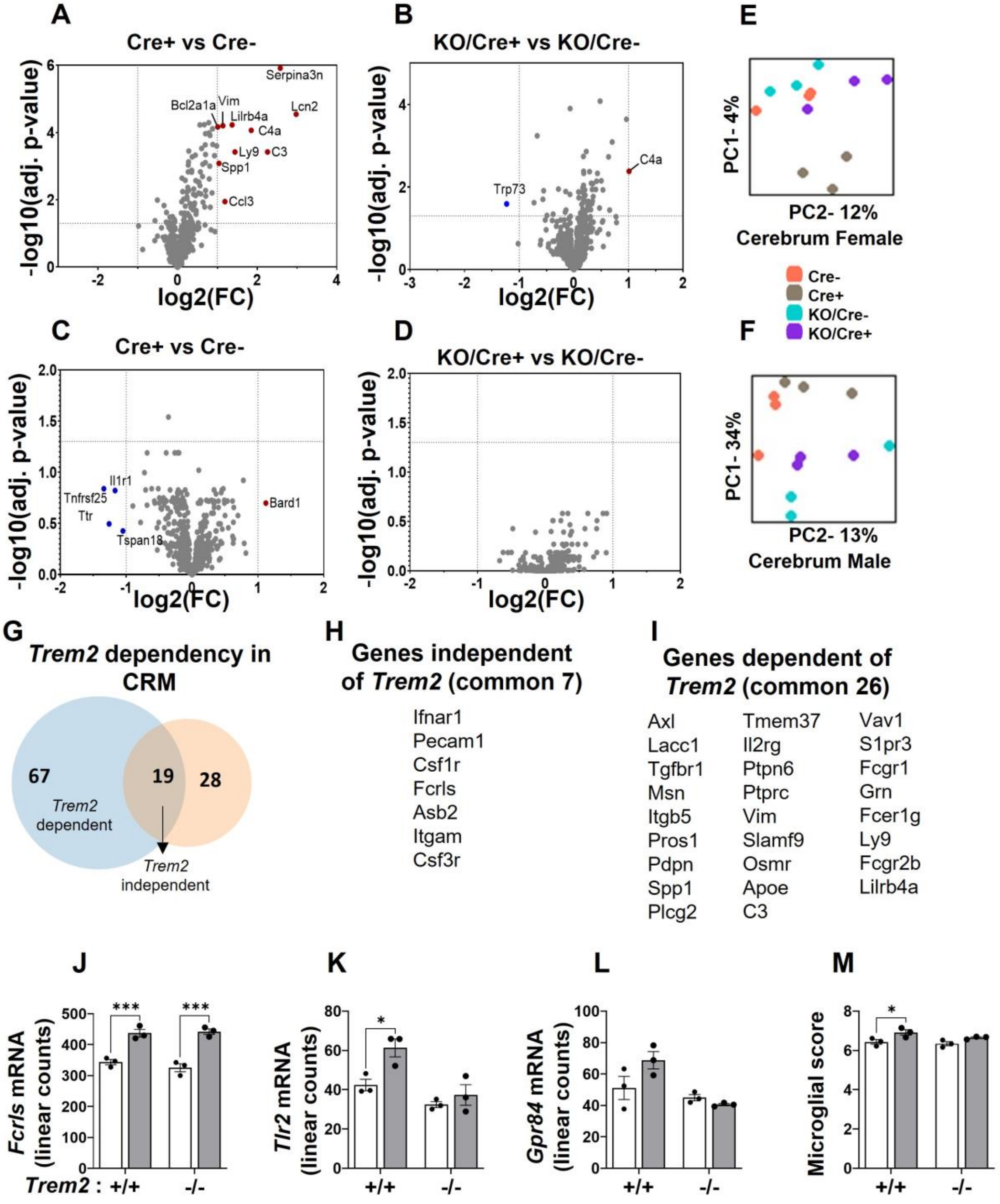


**Fig. S1: The effects of ST deficiency on mouse cerebrum in the presence and absence of *TREM2*.** A-D. Volcano plots showing DEGs from CRM (A. Female Cre+ vs -, B. Female KO/Cre+ vs -, C. Male Cre+ vs -, D. Male KO/Cre+ vs -). E-F. Principal component analysis of mRNA levels obtained from the NanoString Neuroinflammatory Panel of CRM. (E. females, F. males). G. Venn diagram showing overlapping DEGs that are upregulated among genotypes in female CRM (blue: Cre+ vs -) and (orange: KO/Cre+ vs -). H-I. List of DEGs shared between SC and CRM (H. *Trem2* independent manner, I. *Trem2* dependent manner). J. mRNA linear counts of the *Trem2* independent *Fcrls* gene in female CRM (white bar: Cre-, grey bar: Cre+). K-L. mRNA linear counts of genes in female CRM that are *Trem2* dependent (K. *Tlr2*, L. *Gpr84).* M. Microglial cell scores of female CRM based on mRNA levels. Two-way ANOVA with multiple comparisons and a Bonferroni post-hoc test. n = 3. *p < 0.05, **p < 0.01, and ***p < 0.001. Data represents the mean ± S.E.M.

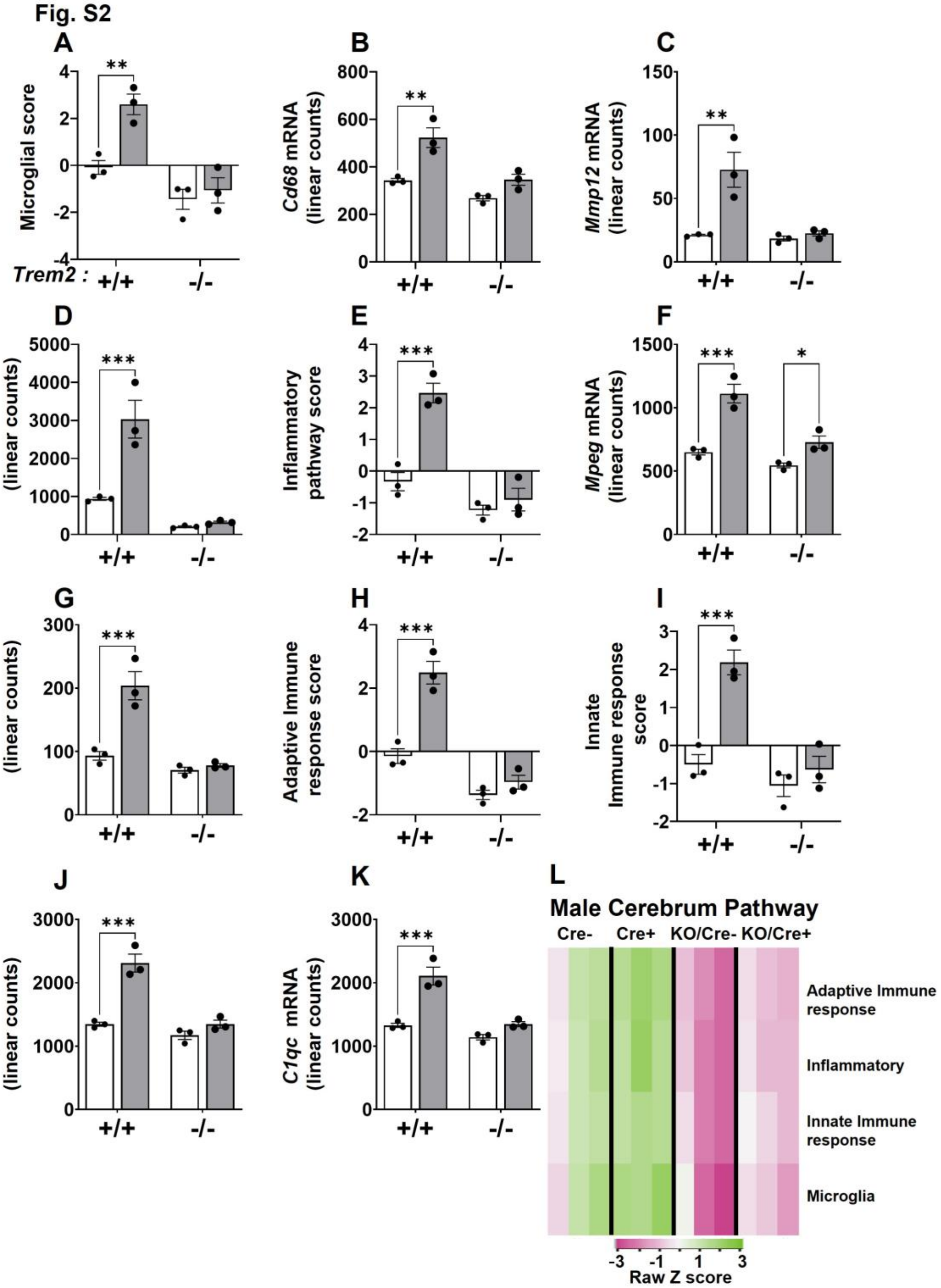


**Fig. S2: *Trem2* regulates ST deficiency-induced microglia-mediated neuroinflammation and immune response pathways in cerebrum.** A. Microglial cell scores of female CRM based on mRNA levels (white bar: Cre-, grey bar: Cre+). B-D. mRNA linear counts of genes in female CRM (B. *Cd68,* C. *Mmp12,* D. *Trem2*). E. Inflammatory pathway score of female CRM. F-G. mRNA linear counts of genes in female CRM (F. *Lilrb4a,* G. *Mpeg).* H-I. Immune response pathway score in female SC (H.

Adaptive, I. Innate). J-K. mRNA linear counts of genes in female CRM (J. *C1qa,* K. *C1qc*). L. Heatmap representing DEGs regulating the listed pathways in male CRM. Two-way ANOVA with multiple comparisons and a Bonferroni post-hoc test n = 3. *p < 0.05, **p < 0.01, and ***p < 0.001. Data represents the mean ± S.E.M.

Fig. S3

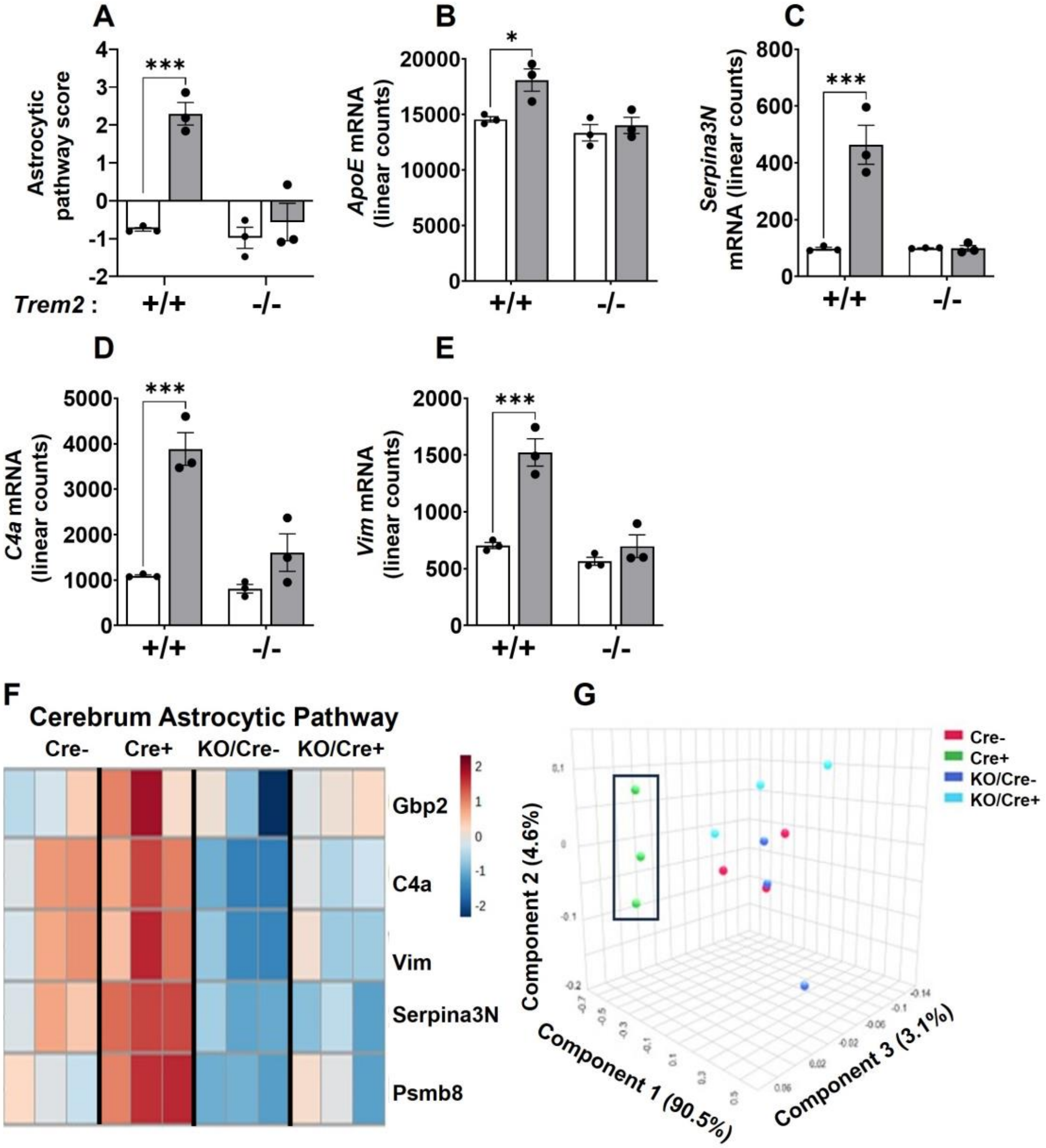


**Fig. S3: *Trem2* regulates ST deficiency-induced activation of astrocytes at the transcriptomic level in cerebrum.** A. Astrocytic score of female CRM based on mRNA levels (white bar: Cre-, grey bar: Cre+). B-E. mRNA linear counts of genes in female CRM (B. *ApoE*, C. *Serpina3N*, D. *C4a*, E. *Vim*). F. Heatmap representing DEGs regulating the astrocytic genes in female CRM. G. PLS-DA analysis of the male CRM. Two-way ANOVA with multiple comparisons and a Bonferroni post-hoc test n = 3. *p < 0.05, **p < 0.01, and ***p < 0.001. Data represents the mean ± S.E.M.

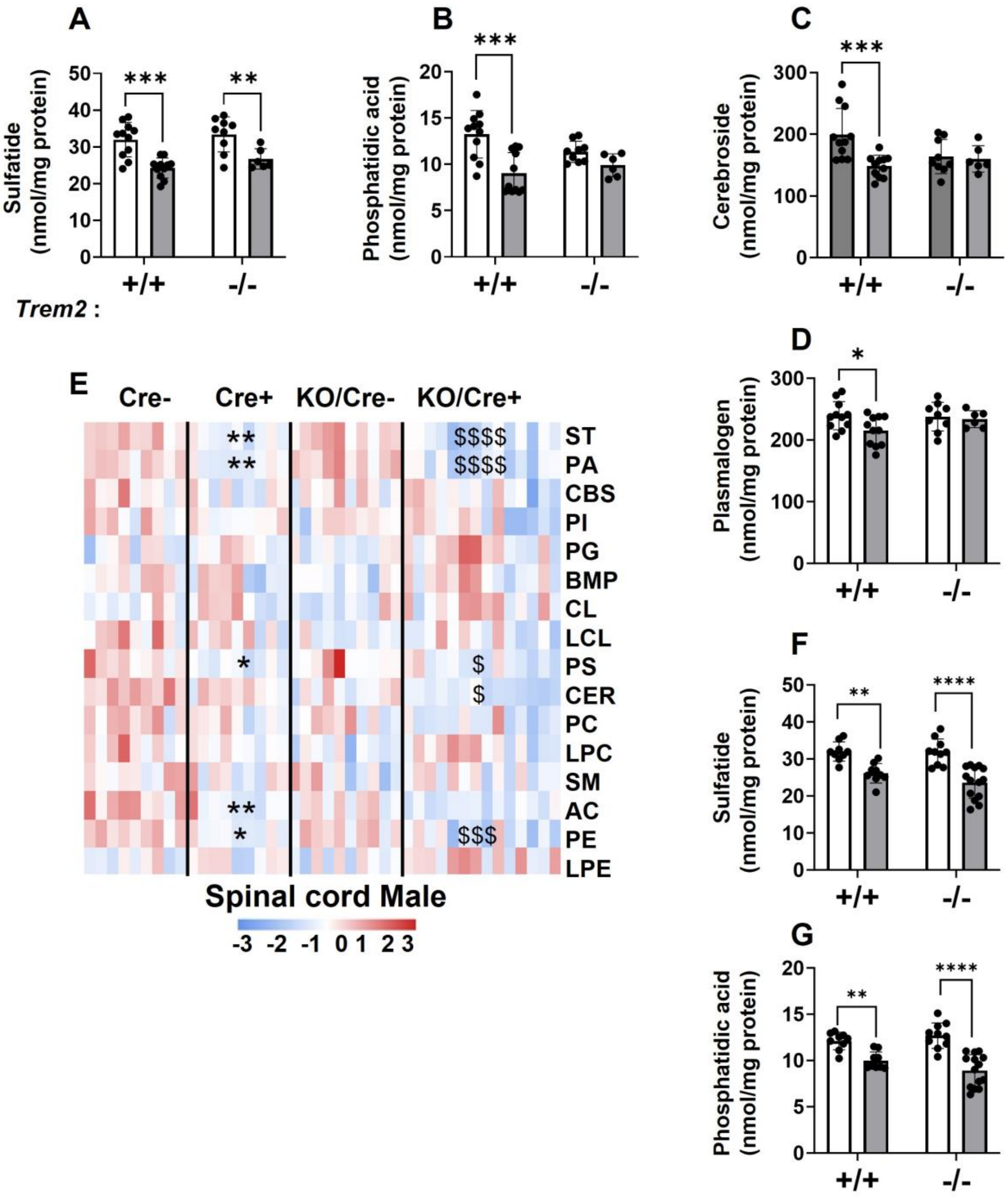


**Fig. S4: Sulfatide deficiency-induced alterations in myelin lipids are independent of *Trem2*.** A-D. Graph (white bar: Cre-, grey bar: Cre+) of each respective lipid in female SC (A. ST, B. PA, C. CBS, D. Plasmalogen). E. Heatmap of total lipid classes from lipidomics data in male SC (* indicates *Cre*+ vs *Cre*- and $ indicates KO/Cre+ vs KO/Cre-). F-G. Graph (white bar: Cre-, grey bar: Cre+) of each respective lipid in male SC (F. ST, G. PA). Two-way ANOVA with multiple comparisons and a Bonferroni post-hoc test n = 5-14. *p < 0.05, **p < 0.01, and ***p < 0.001. Data represents the mean ± S.E.M.

Fig. S5

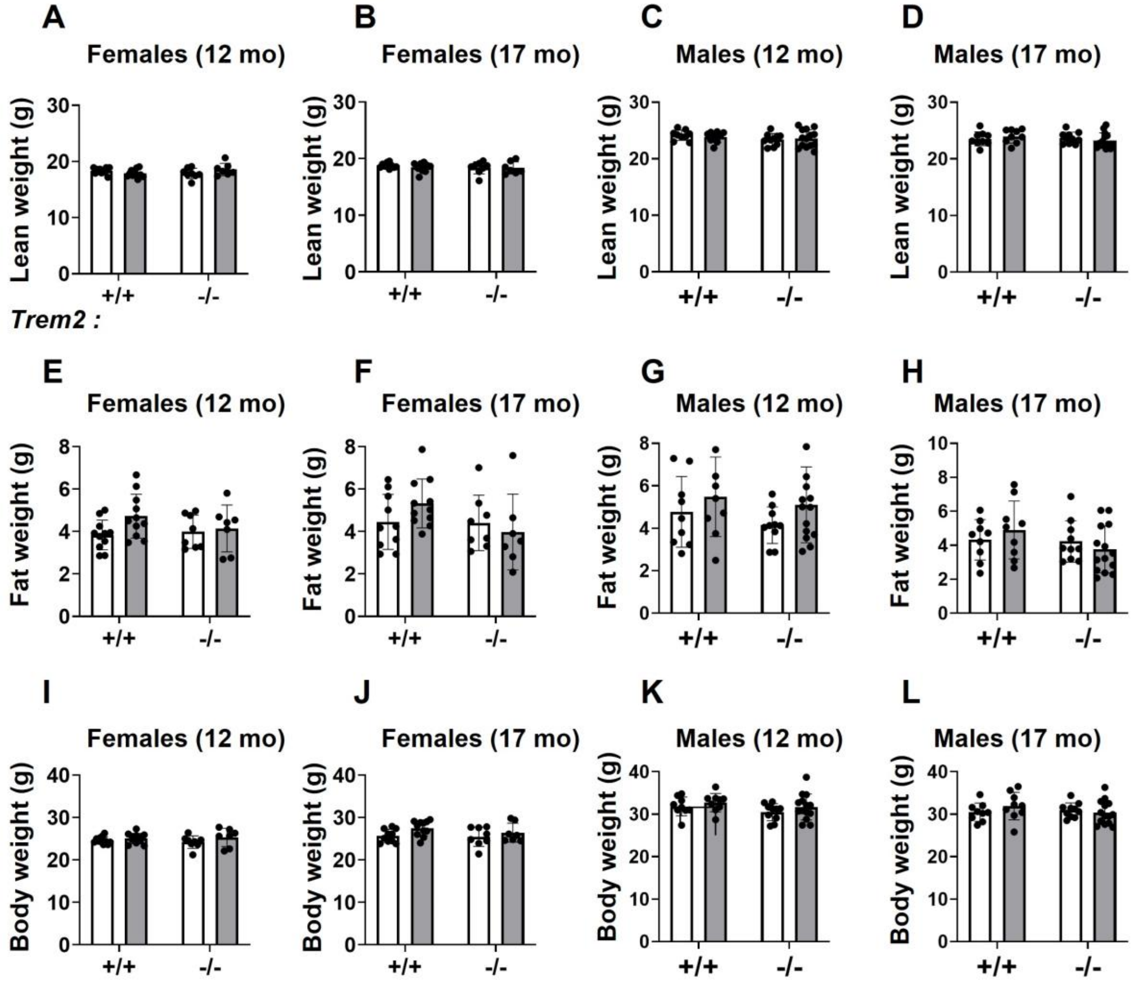


**Fig. S5: Sulfatide deficiency and *Trem2* KO do not affect body composition.** qMRI measurements. A-D. Lean weight (A. 12-mo females, B. 17-mo females, C. 12-mo males, D. 17-mo males). E-H. Fat weight (E. 12-mo females, F. 17-mo females, G. 12-mo males, H. 17-mo males). I-L. Body weight (I. 12-mo females, J. 17-mo females, K. 12-mo males, L. 17-mo males). All graphs (white bar: Cre-, grey bar: Cre+). Two-way ANOVA with multiple comparisons. n = 5-14. *p < 0.05, **p < 0.01, and ***p < 0.001. Data represents the mean ± S.E.M.
